# The novel coronary artery disease risk factor ADAMTS-7 modulates atherosclerotic plaque formation by degradation of TIMP-1

**DOI:** 10.1101/2023.03.06.531428

**Authors:** M. Amin Sharifi, Michael Wierer, Tan An Dang, Jelena Milic, Aldo Moggio, Nadja Sachs, Moritz von Scheidt, Julia Hinterdobler, Philipp Müller, Julia Werner, Barbara Stiller, Zouhair Aherrahrou, Jeanette Erdmann, Andrea Zaliani, Mira Graettinger, Jeanette Reinshagen, Sheraz Gul, Philip Gribbon, Lars Maegdefessel, Jürgen Bernhagen, Hendrik B. Sager, Matthias Mann, Heribert Schunkert, Thorsten Kessler

## Abstract

**Background:** The *ADAMTS7* locus was genome-wide significantly associated with coronary artery disease (CAD). Lack of the extracellular matrix (ECM) protease ADAMTS-7 was shown to reduce atherosclerotic plaque formation.

**Objective:** To identify molecular mechanisms and downstream targets of ADAMTS-7 mediating risk of atherosclerosis.

**Methods:** Targets of ADAMTS-7 were identified by high-resolution mass spectrometry of atherosclerotic plaques from Apoe-/- and Apoe-/-Adamts7-/- mice. ECM proteins were identified using solubility profiling. Putative targets were validated using immunofluorescence, *in vitro* degradation assays, co-immunoprecipitation, and Förster resonance energy transfer (FRET)-based protein-protein interaction assays. *ADAMTS7* expression was measured in fibrous caps of human carotid artery plaques.

**Results:** In humans, *ADAMTS7* expression was higher in caps of unstable as compared to stable carotid plaques. Compared to Apoe-/- mice, atherosclerotic aortas of Apoe-/- mice lacking Adamts-7 (Apoe-/-Adamts7-/-) contained higher protein levels of tissue inhibitor of metalloproteases 1 (Timp-1). In co-immunoprecipitation experiments, the catalytic domain of ADAMTS-7 bound to TIMP-1, which was degraded in the presence of ADAMTS-7 *in vitro.* ADAMTS-7 reduced the inhibitory capacity of TIMP-1 at its canonical target matrix metalloprotease 9 (MMP-9) As a downstream mechanism, we investigated collagen content in plaques of Apoe-/- and Apoe-/-Adamts7-/- mice after Western diet. Picrosirius red staining of the aortic root revealed less collagen as a readout of higher MMP-9 activity in Apoe-/- as compared to Apoe-/- Adamts7-/- mice. In order to facilitate high-throughput screening for ADAMTS-7 inhibitors with the aim to decrease TIMP-1 degradation, we designed a FRET-based assay targeting the ADAMTS-7 catalytic site.

**Conclusion:** *ADAMTS-7,* which is induced in unstable atherosclerotic plaques, decreases TIMP-1 stability reducing its inhibitory effect on MMP-9, which is known to promote collagen degradation and is likewise genome-wide significantly associated with CAD. Disrupting the interaction of ADAMTS-7 and TIMP-1 might be a strategy to increase collagen content and plaque stability for reduction of atherosclerosis-related events.

## Introduction

Atherosclerosis of the coronary arteries is promoted by several risk factors^1^, such as hypertension, diabetes, hypercholesterolemia, smoking, male gender, as well as increasing age and genetic disposition^2^. In recent years, genome-wide association studies (GWAS) have identified multiple common alleles that underlie the genetic risk^3^. One of the strongest loci associated by GWAS with coronary artery disease (CAD) and myocardial infarction (MI) risk^4–6^ represents the extracellular matrix (ECM) protease ADAMTS-7^7^. In experimental studies, lack of Adamts-7, the murine counterpart, was associated with beneficial vascular remodeling^8^ and reduced atherosclerotic plaque formation^9^. Recently, in an experimental study vaccination against ADAMTS-7 successfully reduced atherosclerotic plaque formation under pro-atherogenic conditions and prevented neointima formation and in-stent restenosis^10^. The effects of ADAMTS-7 on vascular remodeling can be explained by degradation of the substrate cartilage oligomeric matrix protein (COMP)^11^, which was also found to mediate ADAMTS-7 related effects in rheumatoid arthritis and represents its first known substrate. Another binding partner of ADAMTS-7 is thrombospondin-1 (THBS1 or TSP1)^8^. In contrast, the precise downstream mechanisms involving ADAMTS-7 in atherosclerotic plaque formation remain incompletely understood.

Here, we observed increased ADAMTS-7 expression in unstable as compared to stable human plaques. In mice, we used a proteome-wide analysis of atherosclerotic aortas to find downstream targets of pro-atherogenic ADAMTS-7. We identified the endogenous inhibitor of metalloproteinases TIMP-1 as a novel target of ADAMTS-7 and verified their interaction at the catalytic site of ADAMTS-7 *in vitro.* As a functional consequence of reduced TIMP-1 levels, increased MMP-9 activity led to reduced plaque collagen content which might influence plaque stability. Finally, we explored the interaction between ADAMTS-7 and TIMP-1 to establish a screening assay for the identification of ADAMTS-7 inhibitors.

## Methods

### Mouse models

Animal experiments were conducted in accordance with the German legislation on protection of animals and approved by the local animal care committee (122–4 (108–9/11)). Mice had ad libitum access to food and water and were housed under a 12 h light-dark cycle. Apoe-/-Adamts7-/- mice were generated by crossbreeding Adamts7-/-^8^ with Apoe^tm1Unc^ (purchased from the Jackson Laboratories, Bar Harbor, USA; subsequently termed Apoe-/-) mice for more than four generations. All experiments were performed on Apoe-/- and Apoe-/- Adamts7-/- mice that were fed a Western Diet (TD88137, Harlan) for either twelve or 16 weeks (specified below) starting at the age of eight weeks for determination of plaque collagen content or high-resolution mass spectrometry of atherosclerotic plaques, respectively.

For analysis of plaque collagen content, male and female mice at a 1:1 ratio were used and sacrificed at an age of 20 weeks (twelve weeks Western Diet). Aortic roots were embedded in optimal cutting temperature compound (Sakura Finetek, Tokyo, Japan) and snap frozen to −80 °C. Frozen samples were cut into 5 μm sections and applied to microscope slides. From the onset of aortic valves, every fifth slide was subjected to tissue staining as described below.

For high-resolution mass spectrometry of atherosclerotic plaques, male animals were analyzed and sacrificed at age 24 weeks (16 weeks Western Diet) by an overdose of pentobarbital (600 mg/kg; Release® 300 mg/ml, WDT, Garbsen, Germany). The arterial tree was perfused through the left ventricle with 20 ml 0.9 % sodium chloride solution. For quantitative detergent solubility profiling aortas were dissected from the ascending part until the bifurcation to the renal arteries and snap frozen in liquid nitrogen and subsequently stored at −80 °C. Sample preparation for proteome analysis and computational mass spectrometry data analysis were performed as described previously^12^. Of note, the data of the Apoe-/- control group were used as cohort #2 in our previous study^12^.

### Cloning of constructs and transfection of eukaryotic cell lines

Different constructs were cloned with the Gateway cloning system (ThermoFisher Scientific, Waltham, USA). A clone containing the consensus *ADAMTS7* coding sequence (NM_014272 Human Untagged Clone) was purchased from Origene (Rockville, USA). To generate Gateway-compatible constructs, the *ADAMTS7* coding sequence was flanked with attB sites by PCR as recommended by the manufacturer. The primers used are listed in **Supplemental Table S1**. A BP reaction was performed to insert the attB site-flanked PCR product into a pDONR221 (ThermoFisher Scientific) entry clone (pDONR221_ADAMTS7). Secondary to plasmid transformation of competent bacteria (New England Biolabs, Ipswich, USA), selection of clones, and plasmid purification, an LR reaction was performed to insert the sequence of interest into a mammalian expression vector (pDEST40; ThermoFisher Scientific). *TIMP1* (HsCD00039987) and *MMP9* (HsCD00042477) were purchased in pDONR221 entry vectors from the Harvard Plasmid Repository and were directly utilized in the LR reactions. We used pDEST40 backbones containing different C-terminal tags (V5/His, FLAG, HA) as indicated in **Supplemental Table S2**. After plasmid transformation of competent bacteria, selection of clones, and endotoxin-free plasmid purification, sequencing on both strands was performed. Secondary to sequence verification, plasmids were used to transfect eukaryotic cell lines as described below. Cells were used 24-48 h after transfection for subsequent assays.

To overexpress *ADAMTS7* in vascular smooth muscle cells (VSMCs), we used a lentiviral construct. To that end, we performed an LR reaction with pDONR221_ADAMTS7 and the pLenti6/V5-DEST vector (ThermoFisher Scientific) to generate the pLenti6_ADAMTS7_V5-DEST construct. Lentivirus production was performed according to the manufacturer’s recommendations in HEK293 E cells using Lipofectamine 3000 (ThermoFisher Scientific). Virus-containing supernatant was collected after 24 and 48 h and combined. After centrifugation at 2,000 rpm, the supernatant was filtered through a 45 μm pore-size filter and stored at −80 °C until further use. Cells were harvested five to six days after transduction.

### Immunoblotting

Cells were washed with Dulbecco’s phosphate buffered saline (PBS; Biochrom, Berlin, Germany), resuspended in radioimmunoprecipitation assay (RIPA) buffer (Sigma-Aldrich, St. Louis, Missouri, USA) or non-reducing lysis buffer (NRL buffer: 25 mM Tris, 150 mM NaCl, 5 mM MgCl_2_, 1 mM DTT, 5 % v/v glycerol and 0.5 v/v % NP40), as appropriate. Lysates were separated from cellular debris by centrifugation. Before loading onto gels, samples were mixed 1:1 with 2x Laemmli buffer (Sigma-Aldrich) and boiled for 5 min at 95 °C. 10 μg of cell lysates were loaded on pre-cast gradient gels (4–20 %, Bio-Rad, Hercules, USA). Proteins were transferred to methanol-activated polyvinylidene difluoride (PVDF) membranes (Bio-Rad) at 100 V for 1,5 h. Membranes were blocked for 1 h at room temperature with 5% dry milk in PBS. The primary antibodies used are listed in **Supplemental Table S3**. Blots were incubated with primary antibodies overnight at 4 °C in 2.5 % dry milk and washed with PBS-T (PBS containing 0.1 % v/v Tween 20, Sigma-Aldrich) before incubation with anti-mouse (#7076S) or anti-rabbit (#7074S) IgG HRP linked antibodies (Cell Signaling Technology, Danvers, USA) at 1:10,000 and 1:100,000 dilutions, respectively, for 1 h at room temperature. After another washing step with PBS-T, bands were visualized by enhanced chemiluminescence (GE Healthcare Life Sciences, Freiburg, Germany). Blots were analyzed using the ImageQuant 800 imaging system (Amersham Biosciences, Amersham, UK) and quantification was performed using ImageJ^13^.

### Immunohistochemistry and immunofluorescence

For immunohistochemistry, sections were fixed with ice-cold 100 % acetone for 10 min at −20 °C and washed with PBS-T (0.1 % v/v Tween 20, Sigma-Aldrich) before blocking with 5 % BSA in PBS (m/v) for 1h at room temperature. After washing, sections were further blocked with Dako REAL Peroxidase-Blocking solution (Agilent, Santa Clara, USA) for 20 min. Next, sections were incubated overnight with primary antibodies (**Suppl. Table S3**) at 4 °C. After washing, biotinylated IgG (1:1000 in 5 % BSA) was added and incubated for 1h at room temperature. 50 % glycerol was then added to each specimen and covered with coverslips.

For immunofluorescence, aortic root sections were rinsed with PBS and fixed in 4 % v/v fresh paraformaldehyde (PFA) at room temperature for 10 min. Next, sections were permeabilized by incubation in PBS containing 0.5 % v/v Triton (x100) for 10 min at room temperature. After washing with PBS, blocking was performed for 1 h at room temperature using PBS containing 5 % m/v BSA and 0.1 % v/v Triton. Sections were incubated with 1:300 dilutions of rabbit anti-Adamts-7 (#201083; Abcam, Cambridge, UK) and goat anti-Timp-1 (#cnl0320021; R&D Systems, Minneapolis, USA) as primary antibodies overnight at 4 °C. Secondary anti-rabbit (#ab150073) and anti-goat (#ab175664; both Abcam) antibodies were added after washing with PBS. Finally, sections were mounted with DAPI mounting solution (Carl Roth, Karlsruhe, Germany).

Images were taken with a THUNDER imaging system (Leica, Wetzlar, Germany) or an Axioscan 7 slide scanner (Zeiss, Jena, Germany).

### Co-immunoprecipitation

Co-immunoprecipitation (Co-IP) was performed using the Dynabeads Co-IP Kit (ThermoFisher Scientific) according to the manufacturer’s instructions. All precipitations were done with 50 mg of cells and antibody-coupled beads (5 μg IP antibody/mg beads). Negative control IPs were performed using IgG coupled beads (Sigma-Aldrich; 5 μg antibody/mg beads). Approximately 10 μg of IP lysate were used for immunoblotting as described above. The antibodies used for IP are listed in **Supplemental Table S4**.

### Matrix metalloproteinase activity assays

#### Fluorescein conjugate gelatine assay

To measure the activity of collagenase and gelatinase, the supernatant of human coronary artery smooth muscle cells overexpressing *ADAMTS7* or a mock plasmid was used with fluorescein conjugates of gelatine (DQ Gelatine from Pig Skin, ThermoFisher). Supernatants were plated on a flat black 96-well plate (Nunclon, ThermoFisher). 200 μl of serum free supernatant were incubated with 0.1 mg/ml of the substrate overnight at 37 °C and fluorescence intensity was measured using an Infinite M200 PRO microplate reader (Tecan, Maennedorf, Switzerland) according to the manufacturer’s recommendations. Background fluorescence was determined and subtracted from each value.

#### Gel zymography

Supernatant of human coronary artery smooth muscle cells overexpressing *ADAMTS7* or a mock plasmid was used. 1 ml of serum-free supernatant was concentrated using 30 μl Gelatine-agarose beads (Sigma-Aldrich) for 16 h at 4 °C. Beads were then washed with PBS and 20 μl sample buffer. Samples were loaded on pre-cast Novex 10% Zymogram Plus Protein Gels (ThermoFisher, Waltham, USA). Electrophoresis was performed at 125 V for 2 h. Afterwards, gels were incubated with renaturing buffer for 30 min, rinsed with water and subjected to developing buffer for 30 min. The developing buffer was then exchanged and further incubation was performed for 48 h. After incubation, the gel was stained with staining solution for 1 h and destained using destaining solution until areas of gelatinolytic activity appeared as sharp bands on a blue background. Gels were scanned for quantification using an ImageQuant 800 imaging system (Amersham Biosciences).

#### Colorimetric assay

Supernatant of HEK 293E cells overexpressing *ADAMTS7* (or mock plasmid) and *TIMP1* were collected and subjected to concentration using 3K Amicon Ultra-15 filters (Sigma-Aldrich). Recombinant MMP-9 from the Matrix Metalloproteinase-9 Colorimetric Drug Discovery Kit (Enzo Life Sciences, New York, USA) was added to the supernatants. Assays were performed according to the manufacturer’s recommendations. Absorbance was measured using an Infinite M200 PRO microplate reader (Tecan).

### TIMP-1 degradation assays

#### In vitro degradation

HEK 293E cells were transiently transfected with *TIMP1* and *ADAMTS7* constructs. After 48 h, supernatants were collected and concentrated with 3K Amicon Ultra-15 filters (Sigma-Aldrich). Cell lysates were prepared as described above. Same amounts of protein of each sample were loaded and the differences in TIMP-1 levels were analyzed using immunoblotting as described above. Quantification was done using ImageJ as described above.

#### Flow cytometry

HEK 293E cells were transiently transfected with *TIMP1-GFP* and *ADAMTS7* constructs or a mock plasmid. After 48 h, cells were washed with PBS and harvested in PBS containing 0.5 % m/v BSA. Cells were filtered through 5 ml Falcon round-bottom polystyrene test tubes with cell strainer snap caps (Falcon, Vancouver, Canada) and DAPI (1:500) was added as viability dye. The numbers and intensity of GFP-positive cells were measured by flow cytometry using a LSRFortessa (BD Biosciences, Franklin Lakes, USA) flow cytometer. The geometric mean fluorescence intensity (geo MFI) index method was used for quantification.

#### Fluorescence microscopy

HEK 293E cells were transiently transfected with *TIMP1-GFP* and *ADAMTS7* constructs. After 48 h, samples were washed with PBS and fixed in 4% v/v PFA for 10 min at room temperature. Permeabilization was induced with 0.5 % v/v Triton in PBS for 10 min at room temperature. Slides were mounted with ROTI Mount FluorCare DAPI (Carl Roth). Images were taken with 10x magnification using a Stellaris 5 confocal microscope (Leica, Wetzlar, Germany). Analysis was performed by measuring the GFP intensity based on the cell count. To obtain cell counts we used a custom macro in ImageJ. GFP intensity divided by cell count was used to quantify TIMP-1-GFP protein levels for each image.

### Picrosirius red staining

Aortic root sections in OCT from Apoe-/-Adamts7-/- and Apoe-/- mice were brought to room temperature and rinsed with PBS. Nuclei were stained with Weigert’s Hematoxylin Kit (Polysciences, Warrington, USA) for 8 min and washed for 10 min under running tap water. Sections were stained in Direct Red 80 stain (Sigma-Aldrich) for 1 h. Specimens were washed in two preparations of acidified water (1.3 % saturated Picric acid). Slides were dehydrated with three preparations of 100 % ethanol, cleared in xylene, and mounted with resinous medium. Images were taken under bright field and polarisation imaging. Quantification of bright field images was performed using a collagen imaging macro (https://imagej.nih.gov/ij/docs/examples/stained-sections/index.html). Polarisation images were analyzed using an in-house developed macro.

### Analysis of ADAMTS7 expression in fibrous caps of human carotid artery plaques

Laser capture micro-dissection of advanced atherosclerotic carotid artery plaques was described previously^14^. In brief, carotid atherosclerotic lesions from the Munich Vascular Biobank^15^ were subjected to microdissection of the fibrous cap and subsequent RNA isolation. RNA was isolated using the RNeasy Micro Kit (Qiagen) and cDNA was synthesized using the Taqman High-Capacity cDNA Transcription Kit (ThermoFisher). Primer assays for *ADAMTS7* and *RPLP0* as housekeeping gene (Hs00276223 and Hs00420895_gH, respectively; both ThermoFisher) were used to detect differences in gene expression between fibrous caps of stable and unstable plaques according to the histomorphologic AHA classification^16^, as described previously^14^. Baseline characteristics of included patients were described by Eken et al.^15^. *ADAMTS7* mRNA levels were visualized and compared as 2^-ΔCt^ values.

### Homogeneous time-resolved fluorescence resonance energy transfer (FRET)

HEK 293E cells were co-transfected with *ADAMTS7-FLAG* and *TIMP1-HA* constructs. Single transfections of constructs were performed as negative controls. After 24 h, cell lysates were prepared using a NRL buffer and final protein concentrations of 1 μg/μl were used. Fluorophore-conjugated antibodies against HA (anti-HA-d2 mAb, #610HADAA) and FLAG (anti-FLAG-M2-Tb cryptate mAb, #61FG2TLA; both Cisbio, Codolet, France) were used. The FRET signal, i.e., the ratio of emission at 665/620 nm wavelength, was measured after 24 h of incubation with an Infinite F200 PRO filter-based microplate reader (Tecan).

### Statistical analysis

Normal distribution of data was assessed using the Kolmogorov-Smirnov test. Test results and subsequently used statistical tests are displayed in **Suppl. Table S5**. Data were analyzed using two-tailed Student’s unpaired or paired t-test (for normally distributed data) or Mann-Whitney test (for non-normally distributed data), as appropriate and indicated in the respective figure legend (and **Suppl. Table S5**). When comparing more than two groups, (RM) one-way repeated measures ANOVA test followed by an appropriate post-test for multiple comparisons was performed when data were normally distributed. To determine statistical outliers, the two-sided ROUT’s test was used. If outliers were removed from the analysis, it is indicated in the respective figure legend. Sample sizes/numbers of replicates are indicated in the figure legends and visualized in the figures (each symbol represents one animal/biological replicate) and data are displayed as mean and s.e.m. P-values below 0.05, in case of investigating more than two groups after adjustment for multiple testing, were regarded as statistically significant. Statistical analyses were performed using GraphPad Prism version 9 for macOS (GraphPad Software, La Jolla, CA, USA).

## Results

### Proteome-wide analysis of the matrisome of atherosclerotic plaques in Apoe-/- and Apoe-/-Adamts7-/- mice identifies Timp-1 as a novel target

ADAMTS-7, an ECM protease^7^, was recently shown to be involved in mediating atherosclerotic plaque formation via its catalytic domain^17^. To unravel potential targets and substrates in atherosclerotic plaque formation, we performed a proteome-wide analysis of mouse aortae – focusing on ECM proteins – after feeding Apoe-/- and Apoe-/-Adamts7-/- mice with a Western diet for 16 weeks. Using solubility profiling as described previously^12^, we detected 3,530 proteins of which 305 were associated with the ECM (**Suppl. Table S6**). The number of detected proteins in both genotypes was comparable (**Suppl. Fig. S1**). Expectedly, known targets of Adamts-7 in vascular remodeling, i.e., thrombospondin-1 (Thbs1)^8^ and cartilage oligomeric matrix protein (Comp)^11^, were detected at numerically higher protein levels in the ECM of Apoe-/-Adamts7-/- compared to Apoe-/- mice (**Fig. 1B**). Further proteins which were detected at higher levels in the absence of Adamts-7 include the tissue inhibitor of metalloproteinases (Timp) 1 (**Fig. 1C**).

**Figure 1:**
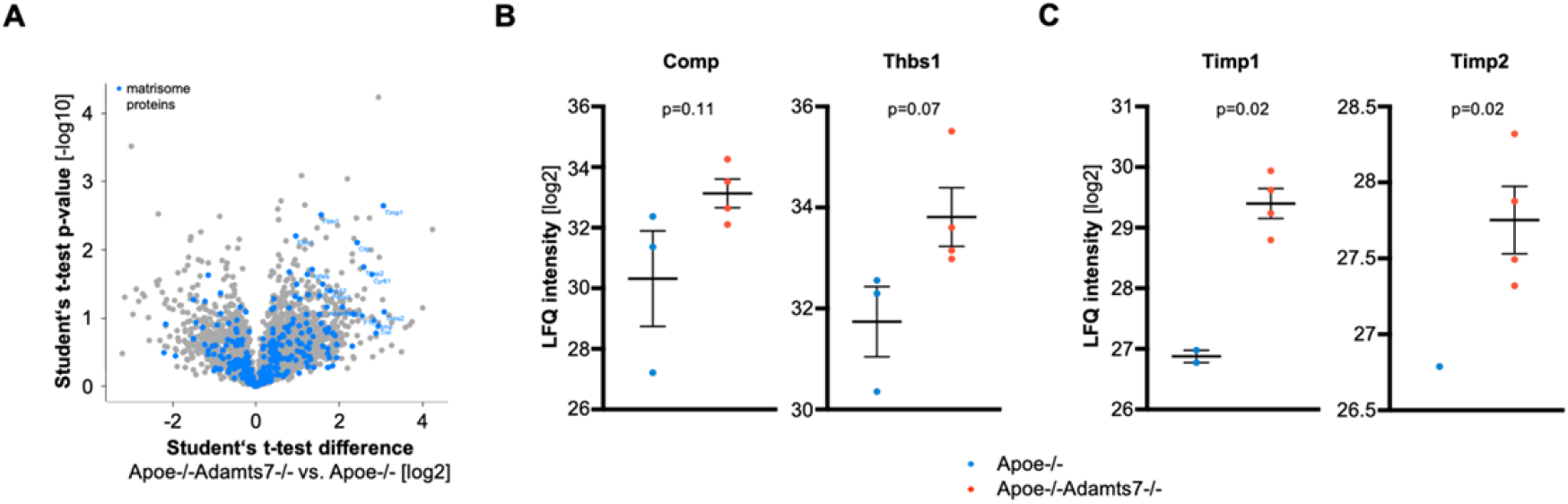
Matrisome profiling of atherosclerotic aortas in Apoe-/- (n=3) and Apoe-/-Adamts7-/- (n=4) mice. **A**. Volcano plot displaying matrisome proteins that were more abundant in Apoe-/-Adamts7-/- mice (right) as compared to Apoe-/- mice. **B**. The known ADAMTS-7 targets Comp and thrombospondin 1 (Thbs1) were detected at lower levels in aortic tissue of Apoe-/- as compared to Apoe-/-Adamts7-/- mice. **C**. As putative novel targets of ADAMTS-7, we observed higher levels for the endogenous tissue inhibitors of MMPs (Timp) Timp-2 and, in particular, Timp-1 (Timp2 was detected in only one of three aortas of Apoe-/- mice). *Student’s t-test with imputation of missing values (i.e., not detected proteins). Each symbol represents one animal. Data are mean and s.e.m*.

### Characterization of the interaction between ADAMTS-7 and TIMP-1

The finding that Timp-1 is detected at higher levels in Apoe-/-Adamts7-/- mice raised the question, whether it is a substrate of Adamts-7 and would directly interact with Adamts-7. To explore this possibility, we first wished to confirm whether Adamts-7 and Timp-1 are co-localized in atherosclerotic plaques. In immunohistochemical stainings, we detected both Adamts-7 and Timp-1 in aortic root plaques of Apoe-/- mice after Western diet for 16 weeks. Immunofluorescence staining suggested co-localization of Adamts-7 and Timp-1 in the plaque region (**Suppl. Fig. S2**). Of note, the knockout of Adamts7 itself did not influence *Timp1* mRNA levels (**Suppl. Fig. S3**). We next sought to investigate whether the proteins directly interact with each other. To that end, ADAMTS7-V5 and TIMP-1-HA constructs were ectopically expressed in HEK 293 cells and protein lysates were subjected to coimmunoprecipitation. To determine the binding domain of TIMP-1 to ADAMTS-7, in addition to the full-length protein, we cloned the N-terminal region containing the catalytic domain and lacking the C-terminal thrombospondin repeats (_ΔTSPr_ADAMTS7-V5) and the C-terminal part containing the disintegrin-like and the THBS1-like domains but lacking the catalytic domain (_Δcat_ADAMTS7-V5) (**Fig. 2A**), and co-expressed these constructs with TIMP-1-HA. Following immunoprecipitation of TIMP-1-HA by an anti-HA antibody, ADAMTS7-V5 was revealed as co-precipitated protein by Western blot, indicating a direct protein-protein interaction (**Fig. 2B**). Importantly, the interaction was also seen when TIMP-1-HA was precipitated from the supernatant (**Suppl. Fig. S4**). The so far known targets of ADAMTS-7, i.e., TSP-1 and COMP, were shown to bind to the C-terminal region of ADAMTS-7 which contains the thrombospondin repeats^8,11^. In further Co-IP experiments, we found an interaction of TIMP-1-HA with the catalytic domain of ADAMTS-7 (**Fig. 2C**), whereas no interaction was detected with the C-terminal part containing the disintegrin-like and the THBS1-like domains (**Fig. 2D**). In summary, these data suggest that TIMP-1 belongs to the small group of ADAMTS-7 interacting proteins that bind to the catalytic domain rather than the C-terminal part of ADAMTS-7.

**Figure 2:**
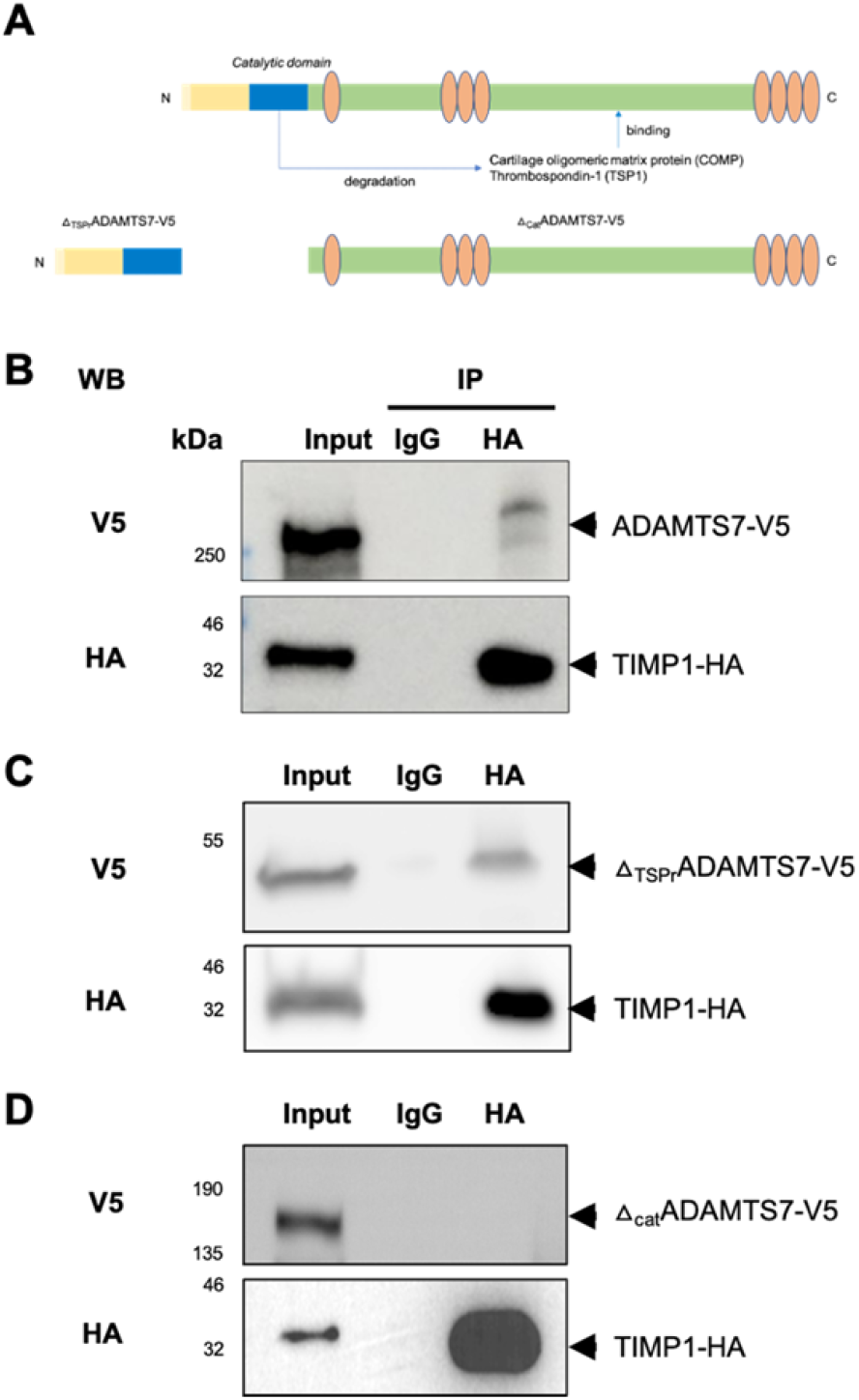
Binding of TIMP-1 to ADAMTS-7. **A**. ADAMTS-7 constructs. We cloned full-length ADAMTS-7 containing a C-terminal V5-tag (ADAMTS7-V5) and two constructs lacking the C-terminal disintegrin domain and thrombospondin repeats (_ΔTSPr_ADAMTS7-V5) or the N-terminal catalytic domain (_Δcat_ADAMTS7-V5), respectively. The constructs were ectopically expressed in HEK 293 cells. **B**. Co-IP of ADAMTS-7 and TIMP-1-HA. After precipitation of TIMP-1-HA, ADAMTS7-V5 was detectable. **C, D**. Binding of TIMP-1-HA to different parts of ADAMTS-7. After precipitation of TIMP-1-HA, _ΔTSPr_ADAMTS7-V5 was detectable (**C**) while in contrast _Δcat_ADAMTS7-V5 was not detectable (**D**) indicating binding of TIMP-1 to the N-terminal part containing the catalytic domain.

### ADAMTS-7 degrades TIMP-1 in vitro

We next sought to investigate whether ADAMTS-7 itself degrades TIMP-1. We therefore performed an *in vitro* degradation assay using HEK 293 cells transiently overexpressing full-length ADAMTS7-V5, the ADAMTS-7 construct lacking the N-terminal region and the catalytic domain (_Δcat_ ADAMTS7-V5), or mock transfection (empty vector). Compared to mock and _Δcat_ADAMTS7-V5 transfection, TIMP-1 protein levels were significantly lower in presence of full-length ADAMTS-7 (ADAMTS7-V5 0.11±0.04 vs. mock: 1.02±0.20 [a.u.], padj=0.02; vs. _Δcat_ADAMTS7-V5: 2.02±0.28 [a.u.], p_adj_<0.01) (**Fig. 3A, B**). We sought to replicate our findings using immunofluorescence and flow cytometry. In line with *in vitro* degradation results, fluorescence microscopy of HEK 293 cells overexpressing TIMP-1-GFP demonstrated lower fluorescence intensity when full-length ADAMTS-7 was present as compared to _Δcat_ADAMTS7-V5 (5.79±0.51 vs. 12.75±0.84 [MFI/cell count], p<0.001; **Fig. 3C, D**). Using flow cytometry, we further analyzed fluorescence intensity in HEK 293 cells overexpressing TIMP-1-GFP in the presence of full-length ADAMTS-7 as compared to _Δcat_ADAMTS7-V5 or mock transfection. We found lower mean fluorescence intensity (MFI) in the presence of full-length ADAMTS-7 (1,034±43.1 vs. mock 2,946±254.2 [MFI], p_adj_=0.01, vs. _Δcat_ADAMTS7-V5 4,764±643.1 [MFI], p_adj_<0.001; **Fig. 3E, F**). Taken together, these data add further evidence that ADAMTS-7 not only binds to TIMP-1 but that TIMP-1 is also subject of degradation by ADAMTS-7.

**Figure 3:**
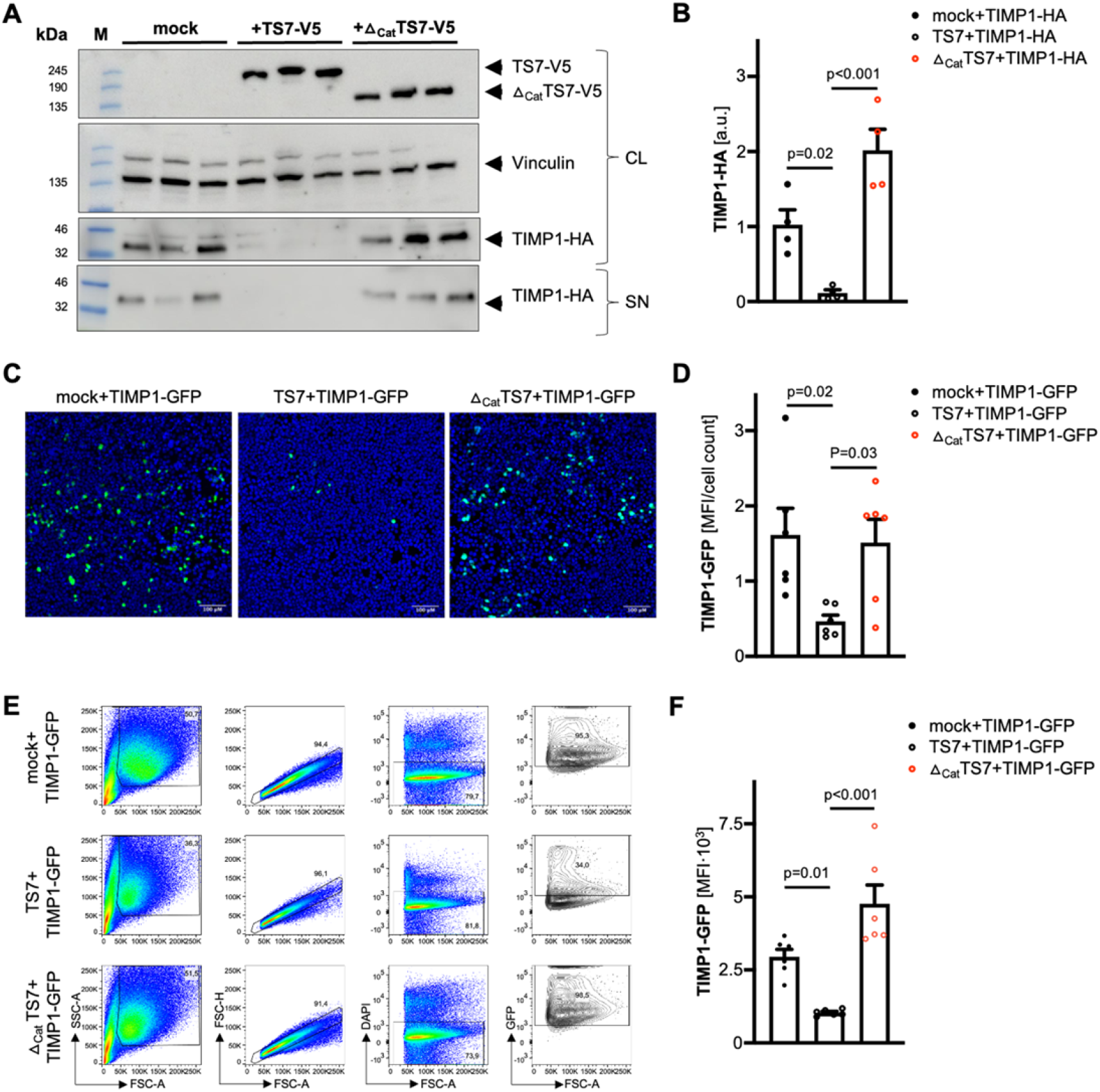
Degradation of TIMP-1 by ADAMTS-7. **A, B**. *In vitro* degradation assay. TIMP-1-HA was either overexpressed with full-length ADAMTS7-V5 (TS7-V5), the C-terminal part lacking the N-terminal catalytic domain (_Δcat_TS7-V5) or an empty vector (mock). ADAMTS-7 constructs and TIMP-1 were detected by immunoblotting in cell lysates (CL) and the supernatant (SN). Vinculin served as housekeeping control. After co-expression with full-length ADAMTS7-V5, less TIMP-1-HA was detectable compared to mock or _Δcat_TS7-V5 co-expression. **C, D**. Fluorescence intensity of TIMP-1-GFP after co-expression with an empty vector (mock), TS7-V5 or _Δcat_TS7-V5 indicates reduced TIMP-1 levels in the presence of full-length ADAMTS-7. **E, F**. GFP-positive cells after overexpression of TIMP-1-GFP with either empty vector (mock), TS7-V5 or _Δcat_TS7-V5. Presence of full-length ADAMTS-7 leads to reduced TIMP-1-GFP fluorescence. *One-way ANOVA with Sidak’s multiple comparisons test. Symbols indicate independent experiments. Data are mean and s.e.m*.

### Effect of ADAMTS-7 and TIMP-1 interaction on MMP-9 activity

After observing scaffolding and degradation of TIMP-1 by ADAMTS-7, we sought to address possible downstream mechanisms. TIMP-1 has been described as an endogenous inhibitor of MMPs, in particular MMP-9, which was associated with plaque instability and adverse cardiovascular events (for an overview see^18^). Importantly, the *MMP9* locus was also identified to be genome-wide significantly associated with CAD risk by GWAS^19^. To examine the influence of ADAMTS-7 on TIMP-1-mediated inhibition of MMP-2/9, human coronary artery smooth muscle cells (hCASMC) were transduced with lentiviral constructs to stably overexpress ADAMTS-7 or a mock control. Activity of endogenous MMP-2/9 in the supernatant was determined using fluorescein-conjugated gelatin substrate. We observed higher activity of MMP-2/9 in supernatants harvested from hCASMC overexpressing ADAMTS-7-V5 in comparison with the mock control (1.22±0.02 vs. 1.08±0.01 [a.u.], p<0.001; **Fig. 4A**). As this assay does not allow individual assessment of MMP-9 and MMP-2 activity, we subsequently performed gel zymography in which gel degradation was measured as a readout of protease activity. Gel zymography revealed higher activity of MMP-9 (ADAMTS-7-V5 205.9±24.7 vs. mock 127.6±8.9 [a.u], p=0.01; **Fig. 4B, C**) and MMP-2 (ADAMTS-7-V5 133.6±8.9 vs. mock 106.2±1.9 [a.u], p=0.01; **Fig. 4B, D**) in samples overexpressing ADAMTS-7 compared to mock showing that ADAMTS-7 reduces the inhibitory capacity of both MMP-2 and MMP-9. To rule out that the observed activities are due to altered endogenous MMP expression, we further performed MMP activity assays using recombinant MMP-9 (rMMP-9). In line with our results investigating endogenous MMP-9 in hCASMC, we found reduced inhibition of rMMP-9 by TIMP-1 in the presence of ADAMTS-7 compared to mock (19.5±0.7 vs. 15.9±1.1 [%], p=0.02; **Fig. 4E**). Of note, ADAMTS-7 alone did not influence MMP-9 activity (**Suppl. Fig. S5**).

**Figure 4:**
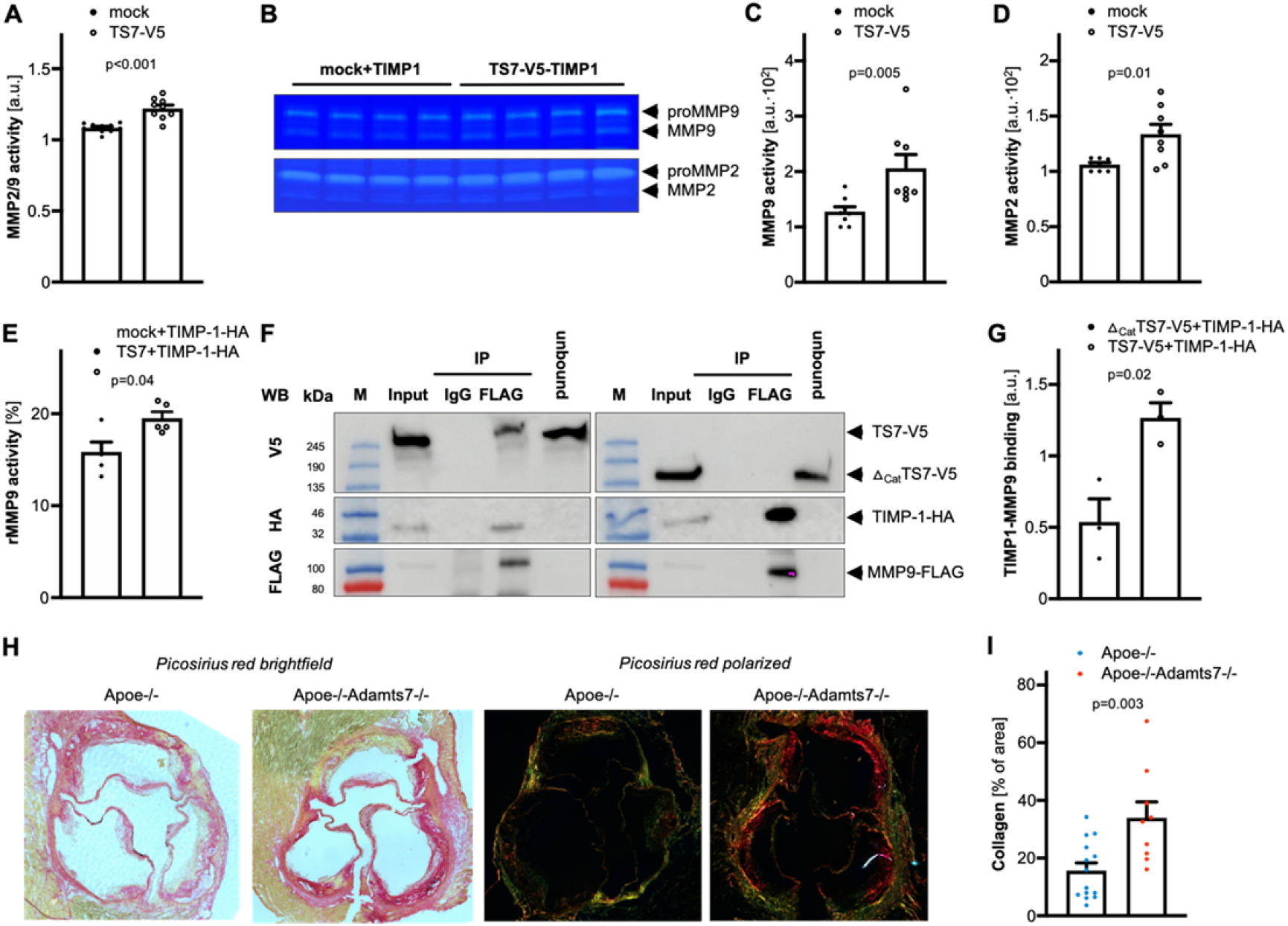
Influence of ADAMTS-7 on inhibition of MMP-9 by TIMP-1. **A**. Endogenous MMP-2-/MMP-9-activity of hCASMC in the presence (TS7-V5) or absence (mock transfection) of full-length ADAMTS-7. Presence of full-length ADAMTS-7 reduced the inhibitory potential of TIMP-1. *Student’s t-test.* **B**-**D**. Gel zymography with endogenous (pro-) MMP-2 and MMP-9 in the presence of either mock or TS7-V5 (**B**). Quantification of MMP-9 and MMP-2 bands reveals higher MMP-9 (**C**) and MMP-2 activity (**D**) in the presence of full-length ADAMTS-7, respectively. *Mann-Whitney (C) and Student’s t-test (D).* **E**. Reduced inhibition of recombinant MMP-9 (rMMP-9) by TIMP-1 in the presence (TS7-V5) as compared to the absence (mock) of full-length ADAMTS-7. *Student’s t-test.* **F, G**. Secondary to precipitating FLAG-tagged MMP-9 (MMP-9-FLAG) more TIMP-1-HA could be recovered in the presence of full-length ADAMTS-7 (TS7-V5) as compared to _Δcat_TS7-V5. *Representative Western Blot. Student’s t-test.* **H, I**. Picosirius red staining of collagen fibers in aortic root sections of Apoe-/- and Apoe-/-Adamts7-/- mice which were fed a Western diet. Mice lacking Adamts-7 displayed increased collagen content as compared to Apoe-/- mice. *Student’s t-test. Symbols indicate independent experiments/animals.*

To further dissect the molecular mechanism of reduced TIMP-1-mediated inhibition of MMP-9, we co-transfected ADAMTS-7 (ADAMTS7-V5 vs. ΔcatADAMTS7-V5), TIMP-1-HA and MMP-9-FLAG in HEK 293 cells. We precipitated MMP-9-FLAG using anti-FLAG-coupled beads and investigated the co-precipitation of the other interaction partners. In the presence of full-length ADAMTS7-V5, less TIMP-1-HA was bound to MMP-9-FLAG as compared to the presence of _Δcat_ADAMTS7-V5 (0.54±0.16 vs. 1.27±0.11 [a.u], p=0.02; **Fig. 4F, G**) suggesting that scaffolding and/or degradation of TIMP-1 by ADAMTS-7 might enhance MMP-9 activity.

Collagen represents a *bona fide* substrate of MMP-9. As a readout of MMP-9 inactivity *in vivo*, we analyzed plaque collagen content in Apoe-/-Adamts7-/- and Apoe-/- mice that had been on a Western diet for 12 weeks. In mice lacking Adamts-7 and displaying increased Timp-1 protein levels in the aortic root (**Suppl. Fig. S6**), collagen content was likewise increased under proatherogenic conditions (33.95±5.5 vs. 16.8±2.6 [%], p=0.004; **Fig. 4H, I**) suggesting that lack of Adamts-7 leads to lower MMP-9 activity.

### ADAMTS-7 expression in stable and unstable atherosclerotic carotid plaques

As depicted in **Fig. 4**, Adamts-7 deficiency was associated with reduced MMP-9 activity and increased plaque collagen content. We therefore next asked whether ADAMTS-7 is associated with plaque stability. To address this, we investigated *ADAMTS7* expression in human atherosclerotic plaques in the carotid artery with a ruptured or stable plaque phenotype in the Munich Vascular Biobank. Specifically, we measured *ADAMTS7* expression in the caps of unstable (n=10; 70% males; age ± s.e.m.: 73.2±1.5 years) and stable plaques (n=10; 70% males; age ± s.e.m.: 73.1±1.9 years) and found that *ADAMTS7* mRNA levels were significantly higher in caps of unstable plaques (stable plaques: median 0.22, interquartile range 0.16-0.63, vs. unstable plaques: median 1.29, interquartile range 0.66-2.25, [2^-ΔCt^], p<0.001; **Fig. 5**).

**Figure 5:**
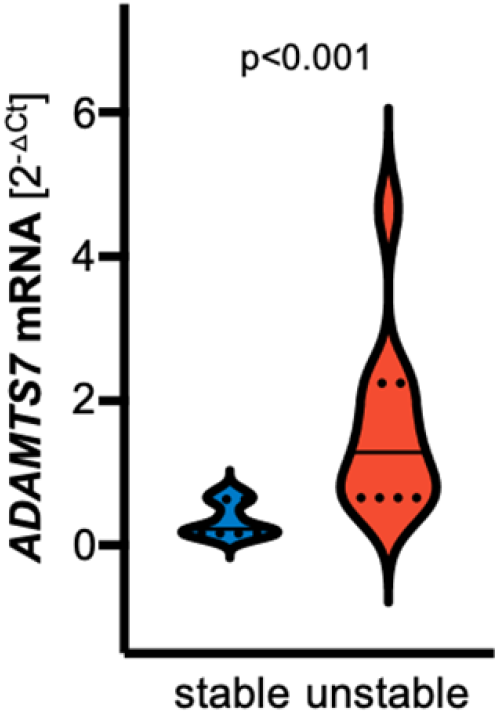
*ADAMTS7* expression in human atherosclerotic carotid plaques. *ADAMTS7* mRNA was detected at higher levels in caps of unstable as compared to caps of stable plaques. *Mann-Whitney test. Symbols indicate independent human individuals.*

### Development of a high-throughput screening-usable protein-protein interaction assay

While the association of ADAMTS-7 with atherosclerosis has been clearly demonstrated in genetic studies in humans^4–6^ and experimental studies in mice^8,9,17,20^, the molecular mechanism remained elusive. Using a proteome-wide analysis of atherosclerotic aorta in Apoe-/- and Apoe-/-Adamts7-/- mice after feeding a Western diet, we identified TIMP-1 as a putative candidate target and substrate. Inhibiting the protein-protein interaction between ADAMTS-7 and TIMP-1 might be a promising therapeutic approach to beneficially influence downstream effects of ADAMTS-7; assays that are suitable for high-throughput screening (HTS) to identify such inhibitors are, however, not available. Therefore, we sought to use time-resolved Förster resonance energy transfer (FRET) to quantify the interaction of ADAMTS-7 and TIMP-1. To that end, we cloned ADAMTS-7-FLAG and TIMP-1-HA constructs. After overexpression in HEK 293 cells, protein lysates were incubated with anti-HA and anti-FLAG antibodies which were bound to the d2 and cryptate fluorophores, respectively. In this assay, we excited ADAMTS-7-FLAG-cryptate at 337 nm wavelength and measured the emission of cryptate at 620 nm and d2 secondary to FRET and 655 nm wavelength. The ratio between 665 nm and 620 nm (FRET ratio) was calculated as the primary readout. While there was no FRET signal detectable in the presence of ADAMTS-7-FLAG or TIMP-1-HA alone, the combination of ADAMTS-7-FLAG and TIMP-1-HA enabled us to detect a strong and reproducible FRET signal (ADAMTS-7-FLAG+TIMP-1-HA 5,484±302.9 vs. mock+TIMP-1-HA 369.5±142.2, p<0.001, vs. ADAMTS-7-FLAG+mock 654.3±240.1 [665/620 ratio], p<0.001; **Fig. 6A**). To verify that the FRET signal is specific for the interaction of ADAMTS-7 and TIMP-1 and to gain first insights into the feasibility as a screening assay for inhibitors, we performed a competition assay of ADAMTS-7-FLAG and TIMP-1-HA with high concentrations of untagged, recombinant TIMP-1. We found a dose-dependent decrease of the FRET signal in the presence of untagged TIMP-1 (untagged TIMP-1: 0 ng 4,954±197.5 vs. 400 ng 4,422±303.2 vs. 800 ng 3,571±143.8 [665/620 ratio], p for trend<0.001; **Fig. 6B**), which is indicative of suitability to search for peptide or small molecule inhibitors. Further confirmation was done by co-expressing non-catalytic domain of ADAMTS-7 with TIMP-1. As expected from the Co-IP results, no FRET signal indicating interaction was detected from overexpression of the non-catalytic domain of ADAMTS-7 with TIMP-1 (**Suppl. Fig. S7**).

**Figure 6:**
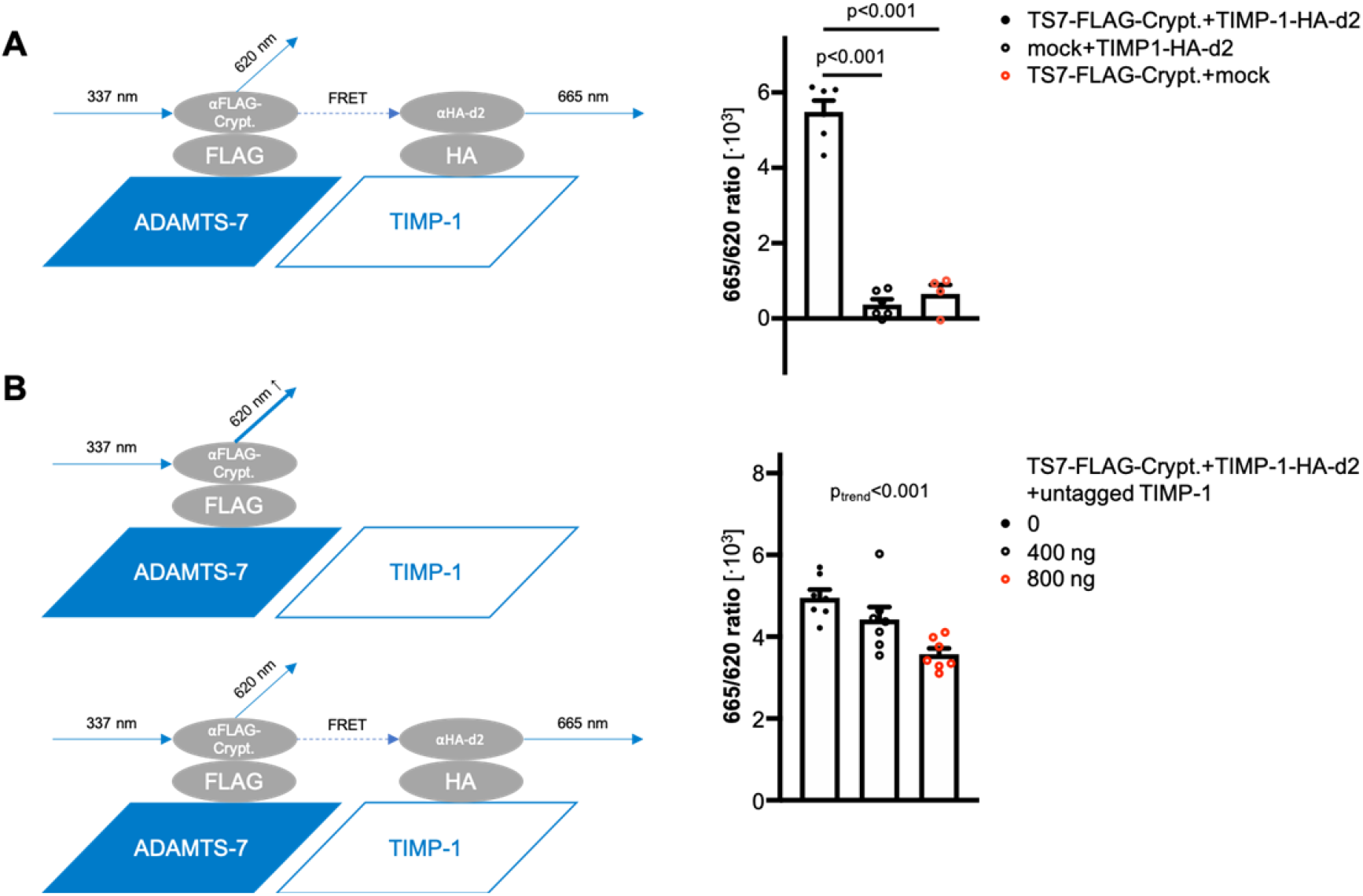
ADAMTS-7-TIMP-1 screening assay. **A**. *Left:* After binding of ADAMTS-7 and TIMP-1 and excitation of cryptate linked to ADAMTS-7, FRET takes place between cryptate and d2. The ratio between emission of d2 and cryptate is used as the readout for interaction between ADAMTS-7 and TIMP-1. *Right:* 665/620 nm ratios for ADAMTS-7 and TIMP-1 and the negative controls, i.e. TIMP-1 or ADAMTS-7 alone. *One-way ANOVA with Sidak’s multiple comparisons test.* **B**. Competition assay. *Left:* In the presence of untagged TIMP-1, emission of cryptate at 620 nm wavelength increases and FRET-induced emission of d2 at 665 nm wavelength decreases. *Right*: 665/620 nm ratios for ADAMTS-7 and TIMP-1 in the presence (400 or 800 ng) or absence (0 ng) of untagged TIMP-1. *One-way ANOVA with post-test for linear trend. Symbols indicate independent experiments*

Taken together, these results replicate the protein-protein interaction of ADAMTS-7 and TIMP-1 using a different, less artificial type of assay. As the binding of TIMP-1 to ADAMTS-7 is a prerequisite for scaffolding and degradation, this assay is suitable for screening against libraries of compounds to identify ADAMTS-7 inhibitors.

## Discussion

ADAMTS-7 represents a novel CAD risk factor but its functional role in atherosclerosis and CAD remains incompletely understood. In this study, we performed a proteome-wide analysis of atherosclerotic aorta tissue from mice on a proatherogenic background which were fed a Western diet. Comparing Apoe-/- and Apoe-/-Adamts7-/- mice, we sought to identify proteomic differences in the presence and absence of Adamts-7. Specifically, bearing in mind the canonical role of ADAMTS-7 as an ECM protease, we focused on the ECM proteome using mass spectrometry analysis of tissue samples after solubility profiling as we described earlier^12^. In mice lacking Adamts-7, the known targets Comp and Thbs1 were detected at higher abundance which confirms previous observations from mechanistic studies in vascular remodeling^8,11^. In addition, we identified TIMP-1, an endogenous inhibitor of metalloproteinases^21^, to be detected at higher protein but not mRNA levels in aortic tissue from mice lacking Adamts-7. The family of TIMPs in humans consists of the four members TIMP-1, TIMP-2, TIMP3, and TIMP-4 (for an overview, see^21^). The different members were shown to inhibit endogenous proteases with different affinities. While all TIMP2-4 were described to inhibit all or most endogenous MMPs, the spectrum of TIMP-1 is rather narrow; in particular, it seems to strongly interact with MMP-9 as it also binds pro-MMP-9. Importantly, considering other proteases particularly from the disintegrin family, TIMP-1 is only known to bind ADAM-10 while TIMP-3 in contrast inhibits a wide range of ADAM and ADAMTS proteases^22^. Since a biological interaction between ADAMTS-7 and TIMP-1 was not known before, we next aimed to verify that both proteins directly interact with each other. Using co-immunoprecipitation, we found that ADAMTS-7 and TIMP-1 directly bind to each other. Importantly, we were not able to demonstrate binding of TIMP-1 to the part of ADAMTS-7 containing the thrombospondin repeats. This is notable as the so far identified targets, including COMP and Thbs1 were shown to bind there^8,11^. In contrast, in this study TIMP-1 bound to the part of ADAMTS-7 containing the catalytic domain. This observation points to an important role of the catalytic domain in atherosclerosis which was recently also demonstrated in a study in which the deletion of only this part of ADAMTS-7 was sufficient to reduce atherosclerotic plaque formation^17^. Interestingly, a previous *in vitro* study showed that ADAMTS-7’s protease function was not affected by TIMP-1, i.e., TIMP-1 itself does not inhibit ADAMTS-7^23^. Taken together, this renders an interaction between the catalytic domain of ADAMTS-7 and TIMP-1 likely to be involved in mediating risk of atherosclerosis.

Although ADAMTS-7 was described as a protease, we considered whether in addition to binding TIMP-1, ADAMTS-7 scaffolds or degrades it or is itself inhibited by TIMP-1. Of note, TIMP-3 has been described to inhibit other ADAM and ADAMTS proteases, e.g. ADAM17^24^ and ADAMTS-2^25^. *In vitro* degradation assays, however, revealed that full-length ADAMTS-7 degrades TIMP-1 while TIMP-1 levels remained stable in the presence of an ADAMTS-7 construct lacking the catalytic domain. Importantly, we observed similar findings using cell-based assays. The consistent in vitro findings along with the observation of increased Timp-1 levels in aorta samples from mice lacking Adamts-7 provide substantial evidence that i) TIMP-1 represents a novel target of ADAMTS-7 and ii) in contrast to TIMP-3, TIMP-1 does not mainly act as an inhibitor of this ADAMTS protease but rather as a substrate.

Previous studies clearly suggested a detrimental role of ADAMTS-7 in vascular biology^8,9,11,17,26^. The role of TIMPs and in particular TIMP-1, in contrast, remains controversial. In a mouse model of atherosclerosis using overexpression of Timp-1 or Timp-2, a study in a model of early atherosclerosis revealed a reduction of atherosclerotic plaques rather for Timp-2 than Timp-1^27^. In line, the same group reported that mice lacking Timp-1 did not develop larger plaques as compared to wild-type mice on a proatherogenic background^28^. This discrepancy might be due to the fact that TIMP-1 is a modulator of a broad range of biological processes. TIMP-1 can exert its effect both in an MMP-dependent and independent manner^29,30^. We here report a reduction of TIMP-1-mediated inhibition of MMP-9 due to scaffolding and degradation by ADAMTS-7 *in vitro*, given that MMP-9 is a *bona fide* interaction partner of TIMP-1^31^. MMP-9 itself is encoded by a CAD risk gene identified by genetic studies^19^ and is one of the main proatherogenic proteases^32^. ECM remodeling caused by MMP-9 activity was shown to be involved in the transition from a stable to an unstable atherosclerotic lesion^18^. In this context, collagen content in the plaque is a readout of MMP-9 activity and we found less collagen content in Apoe-/- as compared to Apoe-/-Adamts7-/- mice. ADAMTS-7 was previously reported to be associated with a vulnerable carotid plaque phenotype^33^. Increased MMP-9 activity, as shown in our study, might contribute to this observation. In line, we found higher expression of *ADAMTS7* in fibrous caps of unstable as compared to stable carotid plaques. In addition, it was shown that changes in ECM equilibrium due to MMP-9 activity induce cell-specific pro-inflammatory or pro-apoptotic activities^34^. As such, MMP-9 promotes proliferation and migration of VSMCs^35^ and proliferation and apoptosis of endothelial cells^21,36^, which contributes to remodeling of the tissue and atherosclerotic angiogenesis. The angiogenesis-inducing potency of pro-MMP-9 is substantially decreased when encumbered by TIMP-1^37^. In addition, TIMP-1 has MMP-9-independent cytokine-like activities^29,30^ and can modulate a broad range of biological processes, e.g. cell growth, proliferation, apoptosis, migration, and angiogenesis, via binding to thus far unknown receptors and thereby inducing specific signaling cascades^38^. Thus, degradation of TIMP-1 by ADAMTS-7 could result in a plethora of further downstream processes regulated by interactions between TIMP-1 and MMP-9.

Taken together, we here provide evidence that the interaction of ADAMTS-7 and TIMP-1 contributes to the modulation of atherosclerotic plaques. As for other CAD risk factors identified by GWAS, there is hope that ADAMTS-7 could be used as a druggable target to prevent and treat atherosclerotic plaque formation. Assays which are able to quantify ADAMTS-7 activity are, however, so far lacking. As binding of TIMP-1 is a prerequisite of both scaffolding and degradation, we used FRET to quantify the protein-protein interaction between ADAMTS-7 and TIMP-1. Importantly, competition with untagged TIMP-1 was able to reduce the signal indicating a potential in screening efforts. This assay forms a sound foundation for it to be screened against relevant small molecule libraries to identify ADAMTS-7 inhibitors as well as modulators of the ADAMTS-7 and TIMP-1 protein-protein interaction.

## Conclusions and limitations

We outlined a molecular mechanism involving the CAD risk gene ADAMTS-7 in atherosclerosis and plaque vulnerability (**Fig. 7**). TIMP-1, being degraded by ADAMTS-7’s catalytic site, represents a novel downstream target that partly explains the role of ADAMTS-7 in CAD. We are aware that ADAMTS-7 might have several further targets contributing to its effects in vascular biology. As such, the roles of COMP and TSP-1 remain to be further investigated. A further CAD risk gene identified by GWAS, sushi, von Willebrand factor type A, EGF and pentraxin domain-containing protein 1 (SVEP1)^39,40^ was shown to be detectable at lower levels after overexpression of ADAMTS-7 before^8^. Current and future research efforts will therefore need to investigate the contribution of other downstream targets and their respective therapeutic potential. To that end, we established a FRET-based protein-protein interaction assay which may be used for high-throughput screening against libraries of small molecules to identify inhibitors that potentially prevent initiation or slow the progression of atherosclerosis. For large-scale screening efforts, limitations such as the preparation of cell lysates and incubation with antibodies need to be overcome. However, the assay can serve as a proof-of-concept for further development. Finally, TIMP-1 as a putative target was identified in mouse aortas which by nature have differences to the human pathophysiology of CAD. The mouse, however, represents an established model in atherosclerosis research and due to the fact that we were able to confirm an interaction between ADAMTS-7 and TIMP-1 using the human proteins renders conservation of this interaction in humans likely.

**Figure 7:**
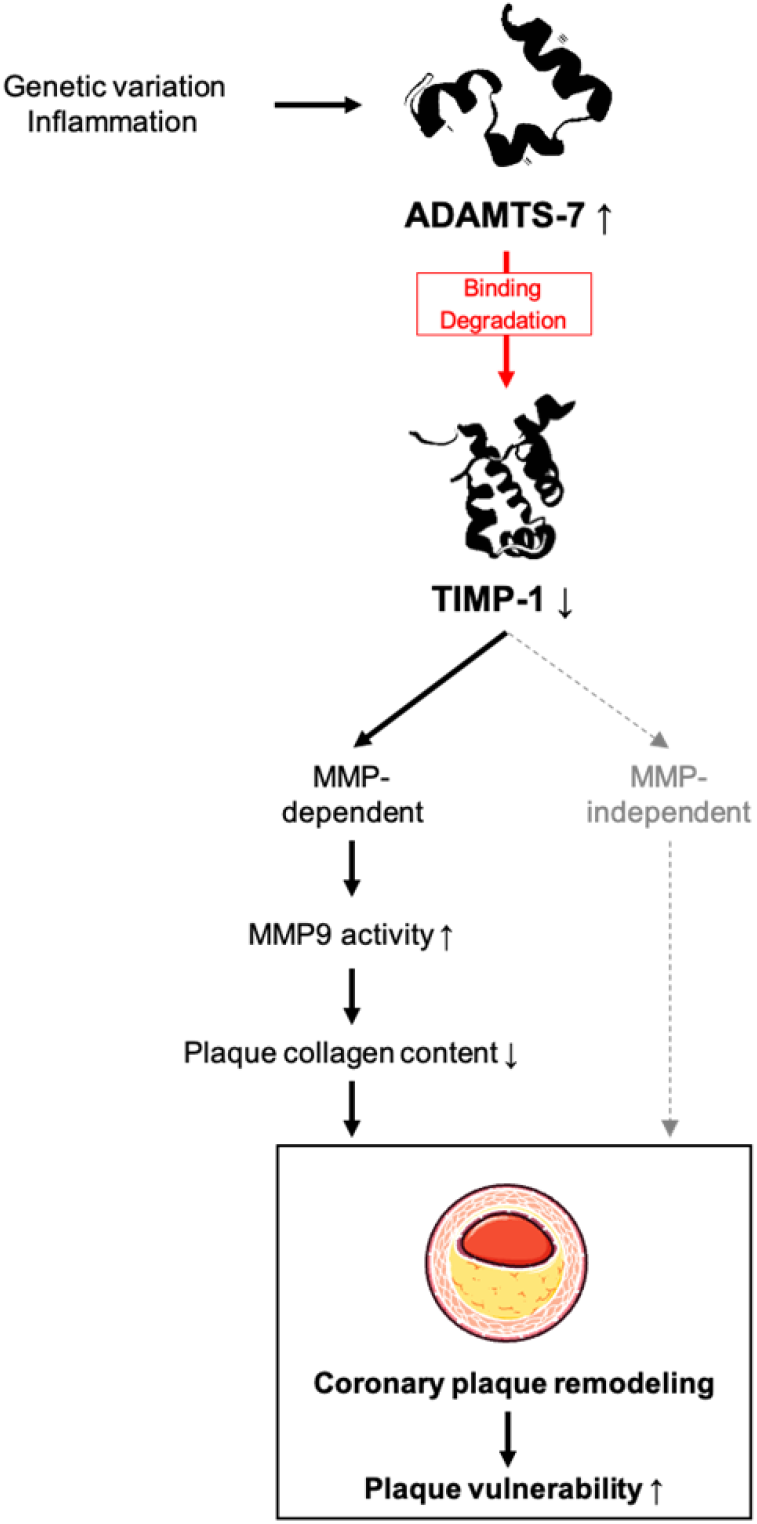
Schematic illustration of the ADAMTS-7/TIMP-1 interaction in atherosclerosis. Genetic variation and inflammation can lead to an upregulation of *ADAMTS7* expression in vascular tissues. As one downstream target, TIMP-1 binds to ADAMTS-7 and is subsequently degraded. Reduced availability of TIMP-1 can lead to MMP-independent and -dependent downstream effects. Among the latter, MMP-9 activity is increased and thereby plaque collagen content reduced. Reduced plaque collagen content can reduce plaque stability and increase plaque vulnerability. Both MMP-independent and -dependent downstream mechanisms may result in altered coronary plaque remodeling and render ADAMTS-7 a promising target in CAD and MI.

## Acknowledgments

The authors thank Christopher Wolf and Sabine Bauer for technical assistance. **Figure 7** contains modified image material available at Servier Medical Art under a Creative Commons Attribution 3.0 Unported License.

## References

1. Virani SS, Alonso A, Aparicio HJ, Benjamin EJ, Bittencourt MS, Callaway CW, Carson AP, Chamberlain AM, Cheng S, Delling FN, Elkind MSV, Evenson KR, Ferguson JF, Gupta DK, Khan SS, Kissela BM, Knutson KL, Lee CD, Lewis TT, Liu J, Loop MS, Lutsey PL, Ma J, Mackey J, Martin SS, Matchar DB, Mussolino ME, Navaneethan SD, Perak AM, Roth GA, Samad Z, Satou GM, Schroeder EB, Shah SH, Shay CM, Stokes A, VanWagner LB, Wang N-Y, Tsao CW, Subcommittee O behalf of the AHAC on E and PSC and SS. Heart Disease and Stroke Statistics—2021 Update: A Report From the American Heart Association. Circulation. 2021;143:e254–e743.

2. Yusuf S, Hawken S, Ounpuu S, Dans T, Avezum A, Lanas F, McQueen M, Budaj A, Pais P, Varigos J, Lisheng L, investigators I study. Effect of potentially modifiable risk factors associated with myocardial infarction in 52 countries (the INTERHEART study): case-control study. Lancet. 2004;364:937–952.

3. Kessler T, Schunkert H. Coronary Artery Disease Genetics Enlightened by Genome-Wide Association Studies. Jacc Basic Transl Sci. 2021;6:610–623.

4. Consortium CADCG. A genome-wide association study in Europeans and South Asians identifies five new loci for coronary artery disease. Nat Genet. 2011;43:339–344.

5. Schunkert H, König IR, Kathiresan S, Reilly MP, Assimes TL, Holm H, Preuss M, Stewart AFR, Barbalic M, Gieger C, Absher D, Aherrahrou Z, Allayee H, Altshuler D, Anand SS, Andersen K, Anderson JL, Ardissino D, Ball SG, Balmforth AJ, Barnes TA, Becker DM, Becker LC, Berger K, Bis JC, Boekholdt SM, Boerwinkle E, Braund PS, Brown MJ, Burnett MS, Buysschaert I, Cardiogenics, Carlquist JF, Chen L, Cichon S, Codd V, Davies RW, Dedoussis G, Dehghan A, Demissie S, Devaney JM, Diemert P, Do R, Doering A, Eifert S, Mokhtari NEE, Ellis SG, Elosua R, Engert JC, Epstein SE, Faire U de, Fischer M, Folsom AR, Freyer J, Gigante B, Girelli D, Gretarsdottir S, Gudnason V, Gulcher JR, Halperin E, Hammond N, Hazen SL, Hofman A, Horne BD, Illig T, Iribarren C, Jones GT, Jukema JW, Kaiser MA, Kaplan LM, Kastelein JJP, Khaw K-T, Knowles JW, Kolovou G, Kong A, Laaksonen R, Lambrechts D, Leander K, Lettre G, Li M, Lieb W, Loley C, Lotery AJ, Mannucci PM, Maouche S, Martinelli N, McKeown PP, Meisinger C, Meitinger T, Melander O, Merlini PA, Mooser V, Morgan T, Mühleisen TW, Muhlestein JB, Münzel T, Musunuru K, Nahrstaedt J, et al. Large-scale association analysis identifies 13 new susceptibility loci for coronary artery disease. Nat Genet. 2011;43:333–338.

6. Reilly MP, Li M, He J, Ferguson JF, Stylianou IM, Mehta NN, Burnett MS, Devaney JM, Knouff CW, Thompson JR, Horne BD, Stewart AFR, Assimes TL, Wild PS, Allayee H, Diemert P, Patel RS, Consortium MIG, Consortium WTCC, Martinelli N, Girelli D, Quyyumi AA, Anderson JL, Erdmann J, Hall AS, Schunkert H, Quertermous T, Blankenberg S, Hazen SL, Roberts R, Kathiresan S, Samani NJ, Epstein SE, Rader DJ. Identification of ADAMTS7 as a novel locus for coronary atherosclerosis and association of ABO with myocardial infarction in the presence of coronary atherosclerosis: two genome-wide association studies. Lancet. 2011;377:383–392.

7. Somerville RPT, Longpré J-M, Apel ED, Lewis RM, Wang LW, Sanes JR, Leduc R, Apte SS. ADAMTS7B, the full-length product of the ADAMTS7 gene, is a chondroitin sulfate proteoglycan containing a mucin domain. J Biol Chem. 2004;279:35159–35175.

8. Kessler T, Zhang L, Liu Z, Yin X, Huang Y, Wang Y, Fu Y, Mayr M, Ge Q, Xu Q, Zhu Y, Wang X, Consortium GMC, Schmidt K, Wit C de, Erdmann J, Schunkert H, Aherrahrou Z, Kong W. ADAMTS-7 inhibits re-endothelialization of injured arteries and promotes vascular remodeling through cleavage of thrombospondin-1. Circulation. 2015;131:1191–201.

9. Bauer RC, Tohyama J, Cui J, Cheng L, Yang J, Zhang X, Ou K, Paschos GK, Zheng XL, Parmacek MS, Rader DJ, Reilly MP. Knockout of Adamts7, a novel coronary artery disease locus in humans, reduces atherosclerosis in mice. Circulation. 2015;131:1202–1213.

10. Ma Z, Mao C, Chen X, Yang S, Qiu Z, Yu B, Jia Y, Wu C, Wang Y, Wang Y, Gu R, Yu F, Yin Y, Wang X, Xu Q, Liu C, Liao Y, Zheng J, Fu Y, Kong W. Peptide Vaccine Against ADAMTS-7 Ameliorates Atherosclerosis and Postinjury Neointima Hyperplasia. Circulation. 2022; epub ahead of print.

11. Wang L, Zheng J, Bai X, Liu B, Liu C-J, Xu Q, Zhu Y, Wang N, Kong W, Wang X. ADAMTS-7 mediates vascular smooth muscle cell migration and neointima formation in balloon-injured rat arteries. Circ Res. 2009;104:688–698.

12. Wierer M, Prestel M, Schiller HB, Yan G, Schaab C, Azghandi S, Werner J, Kessler T, Malik R, Murgia M, Aherrahrou Z, Schunkert H, Dichgans M, Mann M. Compartment-resolved Proteomic Analysis of Mouse Aorta during Atherosclerotic Plaque Formation Reveals Osteoclast-specific Protein Expression. Mol Cell Proteomics. 2017;17:321–334.

13. Schneider CA, Rasband WS, Eliceiri KW. NIH Image to ImageJ: 25 years of image analysis. Nat Methods. 2012;9:671–675.

14. Fasolo F, Jin H, Winski G, Chernogubova E, Pauli J, Winter H, Li DY, Glukha N, Bauer S, Metschl S, Wu Z, Koschinsky ML, Reilly M, Pelisek J, Kempf W, Eckstein H-H, Soehnlein O, Matic L, Hedin U, Bäcklund A, Bergmark C, Paloschi V, Maegdefessel L. Long Non-coding RNA MIAT Controls Advanced Atherosclerotic Lesion Formation and Plaque Destabilization. Circulation. 2021;144:1567–1583.

15. Eken SM, Jin H, Chernogubova E, Li Y, Simon N, Sun C, Korzunowicz G, Busch A, Bäcklund A, Österholm C, Razuvaev A, Renné T, Eckstein H-H, Pelisek J, Eriksson P, Diez MG, Perisic LPM, Schellinger IN, Raaz U, Leeper NJ, Hansson GK, Paulsson-Berne G, Hedin U, Maegdefessel L. MicroRNA-210 Enhances Fibrous Cap Stability in Advanced Atherosclerotic Lesions. Circ Res. 2017;120;633–644.

16. Stary HC, Chandler AB, Dinsmore RE, Fuster V, Glagov S, Insull W, Rosenfeld ME, Schwartz CJ, Wagner WD, Wissler RW. A Definition of Advanced Types of Atherosclerotic Lesions and a Histological Classification of Atherosclerosis: A Report From the Committee on Vascular Lesions of the Council on Arteriosclerosis, American Heart Association. Circulation. 1995;92:1355–1374.

17. Mizoguchi T, MacDonald BT, Bhandary B, Popp NR, Laprise D, Arduini A, Lai D, Zhu QM, Xing Y, Kaushik VK, Kathiresan S, Ellinor PT. Coronary Disease Association With ADAMTS7 Is Due to Protease Activity. Circ Res. 2021;129:458–470.

18. Li T, Li X, Feng Y, Dong G, Wang Y, Yang J. The Role of Matrix Metalloproteinase-9 in Atherosclerotic Plaque Instability. Mediat Inflamm. 2020;2020:3872367.

19. Braenne I, Willenborg C, Tragante V, Kessler T, Zeng L, Reiz B, Kleinecke M, Ameln S von, Willer CJ, Laakso M, Wild PS, Zeller T, Wallentin L, Franks PW, Salomaa V, Dehghan A, Meitinger T, Samani NJ, Asselbergs FW, Erdmann J, Schunkert H. A genomic exploration identifies mechanisms that may explain adverse cardiovascular effects of COX-2 inhibitors. Sci rep. 2017;7:10252.

20. Arroyo AG, Andrés V. ADAMTS7 in cardiovascular disease: from bedside to bench and back again? Circulation. 2015;131:1156–1159.

21. Murphy G. Tissue inhibitors of metalloproteinases. Genome Biol. 2011;12:233–233.

22. Brew K, Nagase H. The tissue inhibitors of metalloproteinases (TIMPs): An ancient family with structural and functional diversity. Biochim Biophys Acta Mol Cell Res. 2010;1803:55–71.

23. Colige A, Monseur C, Crawley JTB, Santamaria S, Groot R de. Proteomic discovery of substrates of the cardiovascular protease ADAMTS7. J Biol Chem. 2019;294:8037–8045.

24. Amour A, Slocombe PM, Webster A, Butler M, Knight CG, Smith BJ, Stephens PE, Shelley C, Hutton M, Knäuper V, Docherty AJP, Murphy G. TNF-α converting enzyme (TACE) is inhibited by TIMP-3. Febs Lett. 1998;435:39–44.

25. Wang W-M, Ge G, Lim NH, Nagase H, Greenspan DS. TIMP-3 inhibits the procollagen N-proteinase ADAMTS-2. Biochem J. 2006;398:515–519.

26. Du Y, Gao C, Liu Z, Wang L, Liu B, He F, Zhang T, Wang Y, Wang X, Xu M, Luo G-Z, Zhu Y, Xu Q, Wang X, Kong W. Upregulation of a Disintegrin and Metalloproteinase With Thrombospondin Motifs-7 by miR-29 Repression Mediates Vascular Smooth Muscle Calcification. Arterioscler Thromb Vasc Biol. 2012;32:2580–2588.

27. Johnson JL, Baker AH, Oka K, Chan L, Newby AC, Jackson CL, George SJ. Suppression of Atherosclerotic Plaque Progression and Instability by Tissue Inhibitor of Metalloproteinase-2. Circulation. 2006;113:2435–2444.

28. Gregoli KD, George SJ, Jackson CL, Newby AC, Johnson JL. Differential effects of tissue inhibitor of metalloproteinase (TIMP)-1 and TIMP-2 on atherosclerosis and monocyte/macrophage invasion. Cardiovasc Res. 2016;109:318–330.

29. Ries C. Cytokine functions of TIMP-1. Cell Mol Life Sci. 2014;71:659–672.

30. Schoeps B, Eckfeld C, Flüter L, Keppler S, Mishra R, Knolle P, Bayerl F, Böttcher J, Hermann CD, Häußler D, Krüger A. Identification of invariant chain CD74 as a functional receptor of tissue inhibitor of metalloproteinases-1 (TIMP-1). J Biol Chem. 2021;297:101072.

31. Olson MW, Gervasi DC, Mobashery S, Fridman R. Kinetic Analysis of the Binding of Human Matrix Metalloproteinase-2 and −9 to Tissue Inhibitor of Metalloproteinase (TIMP)-1 and TIMP-2*. J Biol Chem. 1997;272:29975–29983.

32. Wagsater D, Zhu C, Bjorkegren J, Skogsberg J, Eriksson P. MMP-2 and MMP-9 are prominent matrix metalloproteinases during atherosclerosis development in the Ldlr(-/-)Apob(100/100) mouse. Int J Mol Med. 2011;28:247–53.

33. Bengtsson E, Hultman K, Dunér P, Asciutto G, Almgren P, Orho-Melander M, Melander O, Nilsson J, Hultgårdh-Nilsson A, Gonçalves I. ADAMTS-7 is associated with a high-risk plaque phenotype in human atherosclerosis. Sci rep. 2017;7:3753.

34. Florence JM, Krupa A, Booshehri LM, Allen TC, Kurdowska AK. Metalloproteinase-9 contributes to endothelial dysfunction in atherosclerosis via protease activated receptor-1. PLoS One. 2017;12:e0171427.

35. Dwivedi A, Slater SC, George SJ. MMP-9 and −12 cause N-cadherin shedding and thereby beta-catenin signalling and vascular smooth muscle cell proliferation. Cardiovasc Res. 2009;81:178–86.

36. Bergers G, Brekken R, McMahon G, Vu TH, Itoh T, Tamaki K, Tanzawa K, Thorpe P, Itohara S, Werb Z, Hanahan D. Matrix metalloproteinase-9 triggers the angiogenic switch during carcinogenesis. Nat Cell Biol. 2000;2:737–744.

37. Zajac E, Schweighofer B, Kupriyanova TA, Juncker-Jensen A, Minder P, Quigley JP, Deryugina EI. Angiogenic capacity of M1- and M2-polarized macrophages is determined by the levels of TIMP-1 complexed with their secreted proMMP-9. Blood. 2013;122:4054–67.

38. Ries C. Cytokine functions of TIMP-1. Cell Mol Life Sci. 2014;71:659–72.

39. Investigators MIG and CardiEC, Stitziel NO, Stirrups KE, Masca NGD, Erdmann J, Ferrario PG, König IR, Weeke PE, Webb TR, Auer PL, Schick UM, Lu Y, Zhang H, Dube M-P, Goel A, Farrall M, Peloso GM, Won H-H, Do R, Iperen E van, Kanoni S, Kruppa J, Mahajan A, Scott RA, Willenberg C, Braund PS, Capelleveen JC van, Doney ASF, Donnelly LA, Asselta R, Merlini PA, Duga S, Marziliano N, Denny JC, Shaffer CM, El-Mokhtari NE, Franke A, Gottesman O, Heilmann S, Hengstenberg C, Hoffman P, Holmen OL, Hveem K, Jansson J-H, Jöckel K-H, Kessler T, Kriebel J, Laugwitz KL, Marouli E, Martinelli N, McCarthy MI, Zuydam NRV, Meisinger C, Esko T, Mihailov E, Escher SA, Alvar M, Moebus S, Morris AD, Müller-Nurasyid M, Nikpay M, Olivieri O, Perreault L-PL, AlQarawi A, Robertson NR, Akinsanya KO, Reilly DF, Vogt TF, Yin W, Asselbergs FW, Kooperberg C, Jackson RD, Stahl E, Strauch K, Varga TV, Waldenberger M, Zeng L, Kraja AT, Liu C, Ehret GB, Newton-Cheh C, Chasman DI, Chowdhury R, Ferrario M, Ford I, Jukema JW, Kee F, Kuulasmaa K, Nordestgaard BG, Perola M, Saleheen D, Sattar N, Surendran P, Tregouet D, Young R, Howson JMM, Butterworth AS, Danesh J, et al. Coding Variation in ANGPTL4, LPL, and SVEP1 and the Risk of Coronary Disease. N Engl J Med. 2016;374:1134–44.

40. Winkler MJ, Müller P, Sharifi AM, Wobst J, Winter H, Mokry M, Ma L, Laan SWvan der, Pang S, Miritsch B, Hinterdobler J, Werner J, Stiller B, Güldener U, Webb TR, Asselbergs FW, Björkegren JLM, Maegdefessel L, Schunkert H, Sager HB, Kessler T. Functional investigation of the coronary artery disease gene SVEP1. Basic Res Cardiol. 2020;115:67.

